# Computational modeling of the crosstalk between macrophage polarization and tumor cell plasticity in the tumor microenvironment

**DOI:** 10.1101/430520

**Authors:** Xuefei Li, Mohit Kumar Jolly, Jason T. George, Kenneth J. Pienta, Herbert Levine

## Abstract

Tumor microenvironments contain multiple cell types interacting among one another via different signaling pathways. Furthermore, both cancer cells and different immune cells can display phenotypic plasticity in response to these communicating signals, thereby leading to complex spatiotemporal patterns that can impact therapeutic response. Here, we investigate the crosstalk between cancer cells and macrophages in a tumor microenvironment through *in silico* (computational) co-culture models. In particular, we investigate how macrophages of different polarization (M_1_ vs. M_2_) can interact with epithelial-mesenchymal plasticity of cancer cells, and conversely, how cancer cells exhibiting different phenotypes (epithelial vs. mesenchymal) can influence the polarization of macrophages. Based on interactions documented in the literature, an interaction network of cancer cells and macrophages is constructed. The steady states of the network are then analyzed. Various interactions were removed or added into the constructed-network to test the functions of those interactions. Also, parameters in the mathematical models were varied to explore their effects on the steady states of the network. In general, the interactions between cancer cells and macrophages can give rise to multiple stable steady-states for a given set of parameters and each steady state is stable against perturbations. Importantly, we show that the system can often reach one type of stable steady states where cancer cells go extinct. Our results may help inform efficient therapeutic strategies.

## Introduction

Cancer has been largely considered as a cell-autonomous disease, but recent investigations have highlighted the crucial role of the tumor microenvironment in determining cancer progression (1). Cancer cells can communicate bi-directionally through various mechanical and/or chemical ways with their neighboring cancer cells (2,3), and/or with other components of the tumor microenvironment such as macrophages and fibroblasts, driving aggressive malignancy (4–6). The interconnected feedback loops formed by these interactions can often generate many emergent outcomes for the tumor. Interestingly, many of the latest therapeutic innovations such as immunotherapy are aimed at targeting aspects of the tumor microenvironment instead of the cancer cells (7).

Tumor-associated macrophages (TAMs) are one of the most abundant immune cell populations in the microenvironment (8,9). They have been shown to promote cancer progression in many ways, such as promoting angiogenesis, suppressing function of cytotoxic T lymphocytes, and assisting extravasation of cancer cells (8–12). Generally, the secretome and functions of TAMs have been shown to be close to that of the so-called alternatively activated macrophages (M_2_) (13). In the case of pathogen infections, macrophages that can engulf the pathogen and present processed antigens to adaptive immune cells, are generally characterized as the classically activated ones (M_1_) (14). M_1_ and M_2_ macrophages have different roles during wound healing: while M_1_ macrophages initiate inflammatory responses, M_2_ macrophages contribute to tissue restoration (13). In the context of cancer, M_1_ macrophages have been generally considered anti-tumor (15–17), whereas M_2_ macrophages have been considered as pro-tumor (10).

However, macrophage polarization is not as rigid as the differentiation of T lymphocytes (18); instead, M_1_, M_2_, and any intermediate state(s) of macrophage polarization are quite plastic (13,19,20). Thus, the idea that reverting TAMs in the cancer microenvironment to its cancer-suppressing counterpart is tempting, the proof of principle of which has been demonstrated in mice models (21–25).

Not only TAMs, but also cancer cells themselves can be extremely plastic, a canonical example of which is epithelial-mesenchymal plasticity, i.e. cancer cells can undergo varying degrees of Epithelial-Mesenchymal Transition (EMT) and its reverse Mesenchymal-Epithelial Transition (MET) (26,27). EMT/MET has been associated with metastasis (28), chemoresistance (29), tumor-initiation potential (30), resistance against cell death (31), and evading the immune system (32).

Importantly, macrophages and cancer cells can interact with and influence the behavior of one another, as shown by many *in vitro* experiments. Specifically, some epithelial cancer cells are capable of polarizing monocytes into M_1_-like macrophages (33). Forming a negative feedback loop, these M_1_-like macrophages can decrease the confluency of the cancer cells that polarized them (33). Moreover, pre-polarized M_1_ macrophages can induce senescence and apoptosis in human cancer cell lines A549 (34) and MCF-7 (35). Intriguingly, factors released by apoptosis of cancer cells can convert M_1_ macrophages into M_2_-like macrophages (35), thus switching macrophage population from being tumor-suppressive to being tumor-promoting. On the other hand, mesenchymal cancer cells can polarize monocytes into M_2_-like macrophages (33,36,37), that can in turn assist EMT (37,38). Thus, the interaction network among macrophages and cancer cells is formidably complex, and the emergent dynamics of these interactions can be non-intuitive (39) yet are often crucial in deciding the success of therapeutic strategies targeting cancer and/or immune cells. For example, even if TAMs at some time can be converted exogenously to M_1_-like macrophages, if most cancer cells still tend to polarize monocytes to TAMs, other coordinated perturbations may be needed at different time-points to restrict the aggressiveness of the disease.

Here, we develop mathematical models to capture the abovementioned set of interactions among cancer cells in varying phenotypes (epithelial and mesenchymal) and macrophages of different polarizations (M_1_-like and M_2_-like). We characterize the multiple steady states of the network that can be obtained as a function of different initial conditions and key parameters, and thus analyze various potential compositions of cellular populations in the tumor microenvironment. This *in silico* co-culture system can not only help explain *in vitro* multiple experimental observations and clinical data, but also help acquire novel insights into designing effective therapeutic strategies aimed at cancer cells and/or macrophages.

## Results

### Crosstalk among cancer cells and macrophages can lead to two distinct categories of steady states

We first considered the following interactions in setting up our mathematical model: a) proliferation of epithelial and mesenchymal cells (but not that of monocytes M_0_, or macrophages M_1_ and M_2_), b) EMT promoted by M_2_-like macrophages and MET promoted by M_1_-like macrophages, c) polarization of monocytes (M_0_) to M_1_-like cells aided by epithelial cells, and that to M_2_-like cells aided by mesenchymal cells, d) induction of senescence in epithelial cells by M_1_-like macrophages (Figure 1A). No inter-conversion among M_1_-like and M_2_-like macrophages or cell-death of macrophages is considered here in this model (hereafter referred to as ‘Model I’ see Methods).

Furthermore, this model also considers that mesenchymal cells can secrete soluble factors, such as TGFβ, that can induce or maintain the mesenchymal state in autocrine or paracrine manners (40,41). Therefore, we hypothesized that mesenchymal cells can resist M_1_-promoted MET. However, this resistance might not fully suppress MET, as MET still happens in the presence of M_1_ macrophages (38). Therefore, in Eq. [1], we assume a Hill-like function to represent the resistance to M_1_-promoted MET and the resistance term saturates to a finite value as a function of mesenchymal population M. Similarly, epithelial cells adhere to each other via E-cadherin, sequestering β-catenin on the membrane, thus interfering with the induction of EMT (42). Thus, we hypothesized that the M_2_-promoted EMT can be inhibited by epithelial cells in a cooperative manner, because this inhibition of EMT requires direct physical cell-cell contact and hence involves multiple epithelial cells. Therefore in Eq. [1], we assume a Hill-like function to represent this resistance to M_2_-dependent EMT and the resistance term saturates to a finite value as a function of epithelial population E. Furthermore, the epithelial-cell-dependent term has a cooperativity parameter k to represent the effects of E-cadherin-β-catenin interaction between multiple epithelial cells.

Note that in all of our calculations, we assumed that there is a carrying-capacity of cells (N_max_, including both cancer cells and macrophages) in the co-culture system *in vitro*. We vary the initial number of different cells while keeping the total number of macrophages (M_1_+M_2_+M_0_) to be a constant (M_c_). Therefore, the maximum number of cancer cells will be N_max_-M_c_.

In Model I, the final populations of E, M, M_1_ and M_2_ cells are simply determined by the following equations:

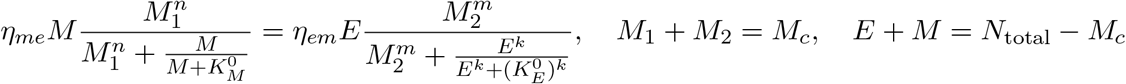

The above equations give two categories of stable steady states: a) state I, dominated by epithelial cancer cells, and b) state II, dominated by mesenchymal cancer cells. The final steady state of the system depends on the number of M_2_ macrophages in that state; since there are only three equations for four unknowns, M_2_ can be used as a parameter specifying (possibly discrete set of) solutions. As soon as M_2_/N_max_ crosses a threshold, the system switches from state I to state II (Figure 1B). This prediction is largely robust to parameter variation (Figure S1).

Next, we explored what factors could change the qualitative behavior of the model, i.e., enable a smoother and continuous transition of epithelial and mesenchymal percentages as a function of the M_2_-macrophage population. We identified that reducing the cooperativity of epithelial cancer cells in their resistance of M_2_-promoted EMT can lead to a loss of the feature with two-types of steady states (Figure 1C, blue, cyan and light-green lines). Conversely, increasing the cooperativity of M_2_-promoted EMT or M_1_-promoted MET can expand the region of overlapping between state I and II of the system (Figure 1C, green lines). Another factor that can alter the behavior of the model is the initial number of monocytes in the system. At high enough number of monocytes, the absolute number of cancer cells will be small, thus the effect of the cooperativity between epithelial cancer cells will be reduced, and consequently, as discussed above, the overlap of the two types of steady states of the system will disappear (Figure 1D). Together, this increased propensity of multi-stability in the system upon including cooperative effects in the interactions among different species (or variables) is reminiscent of observations in models of biochemical networks (43).

Note that using the M_2_/N_max_ as the “control variable” is specific for Model I, because there is no interconversion between M_1_ and M_2_ macrophages. For a given set of parameters, the steady state level of M_2_ can vary continuously from 0 to M_c_ (Figure 1B-D), and in practice is determined by the initial conditions (initial number of E, M, M_1_, M_2_, and M_0_). For models in the following sections, M_2_/N_max_ will be shown to be fixed to discrete allowed values (instead of continuously varying) for a given set of parameters. For example, if we add a very small conversion rate between M_1_ and M_2_, the steady state of the system will collapse to only one steady state (Figure S2): on the shorter time scale, the trajectory of the system (on the phase plane of E and M_2_) will first converge to one steady state in Model I (with the same M_2_/N_max_); on the longer time scale, as determined by the value of inter-conversion rate between M_1_ and M_2_, the system will slowly evolve to the steady state (blue dot in Figure S2), following along the now-approximate steady states in Model I.

**Figure 1.**
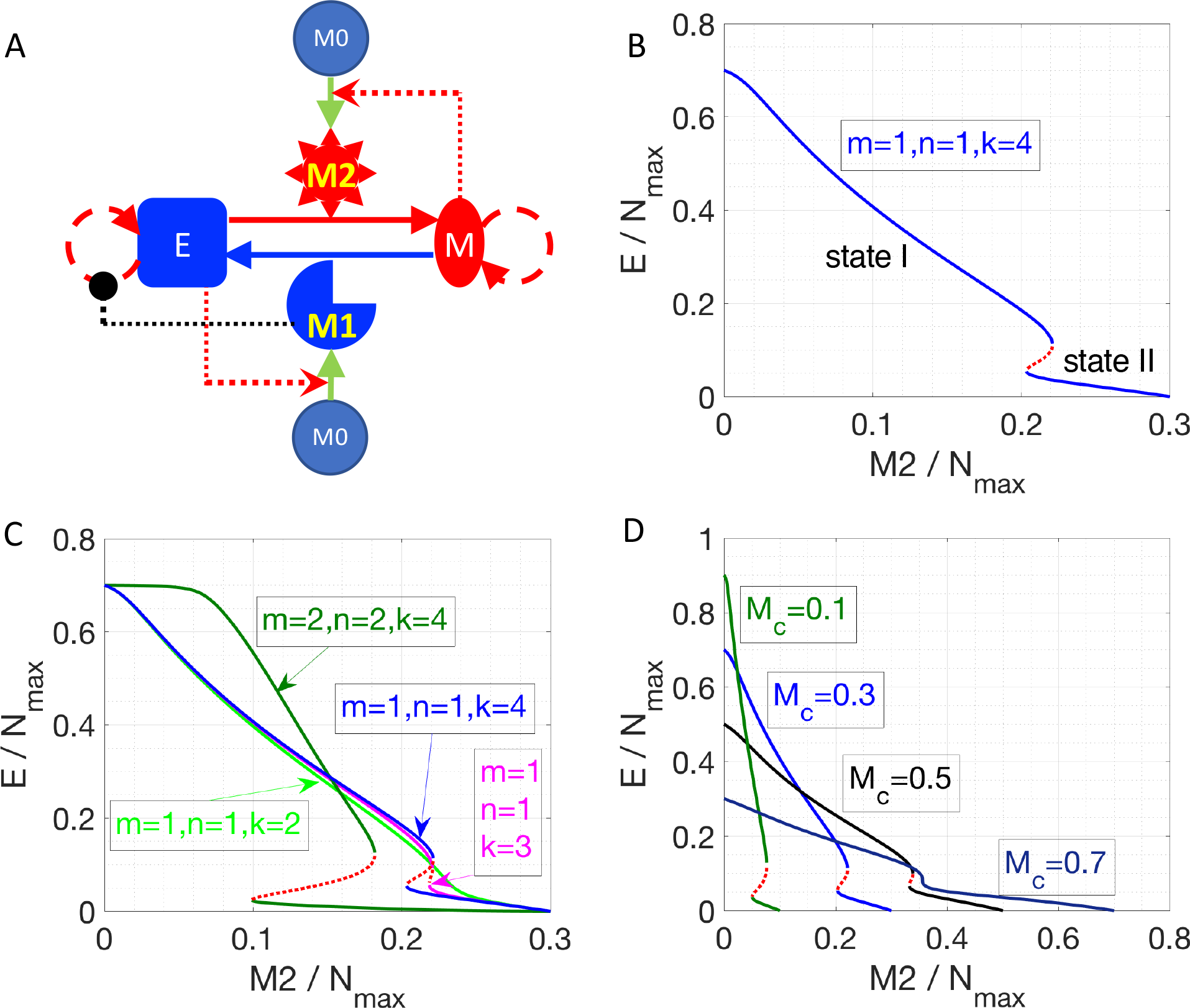
Cancer-immune interplay can give rise to the co-existence of two types of steady states. A. Interaction network for Model I. Conversions between cells are indicated by solid lines. Cell proliferation is indicated by dashed lines. Inhibition (in black) and activation (in red) is indicated by dotted lines. B. Steady states of the epithelial population are plotted as a function of M_2_-like macrophage population. Stable steady states are plotted in solid blue lines and unstable steady states are plotted in dotted red line. The key parameters are as indicated. C. As the cooperativity of epithelial cancer cells in their resistance of M_2_-promoted EMT is reduced, i.e., k=4,3,2 (blue, cyan, and light-green line), the overlapped region between state I and II shrinks and then disappears. Increasing the cooperativity of M_2_-promoted EMT or M_1_-promoted MET, i.e., m and n increase from 1 to 2, can expand the overlapped region (between state I and II) of the system (dark-green lines). Stable steady states are plotted in solid lines and unstable steady states are plotted in dotted lines. D. As the total number of macrophages (M_c_) increases, the overlapped region (between state I and II) of the system shrinks and then disappears.

### Cancer-cell enhanced interconversion between M_1_ and M_2_-like macrophages lead to bi-stability

In the next iteration of our model (hereafter referred to as ‘Model II’), we included the possibility of interconversion between M_1_-like and M_2_-like macrophages, as reported in the literature (13,35,44,45) (Figure 2A, see Methods). We first assume constant interconversion between M_1_ and M_2_ (with rates denoted as η^0^_21_ and η^0^_12_) and investigate the effects of varying the rate of conversion of mesenchymal cells to epithelial cells (η_me_). At small values of η_me_, the system has small number of epithelial cells, which is equivalent to state II in Model I; with increasing η_me_, a threshold is crossed, and the system can switch to states with larger number of epithelial cells, which is equivalent to state I in Model I (solid black lines, Figure 2B). Thus, we can observe bi-stability of cancer cell population in this system. However, the populations of macrophages stay constant as a function of η_me_, since the M_1_/M_2_ ratio is simply determined by η^0^_21_/η^0^_12_ according to Eq. [2] when η_12_=η_21_=0). In this model, we focused on increased cooperativity of M_2_-assisted EMT and M_1_-assisted MET (i.e., m=n=2), because the interconversion between M_1_ and M_2_, in absence of such cooperativity (as considered in model I with m=n=1; Fig 1B), gives rise to a narrower bi-stable region (Figure S3). Note again that in our calculations, we assumed that there is a carrying-capacity of cells (N_max_, including both cancer cells and macrophages) in the co-culture system *in vitro*. Again the total number of macrophages is constant, since no cell death or cell division of macrophages is considered.

We chose η_me_ as a control parameter, because it can act as a bottleneck for the transition between the two types of stable steady states of the system. Lowering the transition threshold of η_me_ can be helpful in a sense that the system can potentially stay at state I (high epithelial state, supposed to be less aggressive) only. Due to the inherent symmetry in the network, the effect of lowering the threshold of η_me_ can be recapitulated via other perturbations, such as a smaller rate of M_1_-assisted MET (η_em_), lower M_1_ to M_2_ conversion rate (η_12_), or higher M_2_ to M_1_ conversion rate (η_21_) (Figure S4).

We next consider the case where interconversion between M_1_-like and M_2_-like macrophages is enhanced by cancer cells, i.e., mesenchymal cells enhance the M_1_-to-M_2_ transition, while epithelial cells enhance M_2_-to-M_1_ transition. In this case, we observe the existence of a bi-stable region, and the range of the epithelial-cell-low solution (equivalent to state II) is wider than that for the previous case without any effects of epithelial (E) and mesenchymal (M) cancer cells on M_1_-M_2_ interconversion (Figure 2B and 2C, solid blue lines). The reason for the expanded epithelial-cell-low region is that the positive feedback loop between mesenchymal and M_2_ cells makes the mesenchymal-and M_2_-dominated state more stable. Therefore, a higher η_me_ is required to compensate for the effects of low M_1_ population. For therapeutic purposes, state I is favored, for it is believed that epithelial cells are typically less aggressive than mesenchymal ones. Therefore, symmetrical mutual enhancement might not be helpful for the therapy because of the expanded epithelial-low (mesenchymal-high) region. For the same reason, the lower threshold of η_me_ to switch from state II to state I also shift to a higher value, which means that for a small η_me_, the system will only have one stable steady state, i.e., state II with high number of mesenchymal cells.

In order to drive the system to be bi-stable at a smaller η_me_, we tested the case with asymmetrical interconversion rates between M_1_ and M_2_, i.e. higher conversion rate from M_2_ to M_1_ enhanced by epithelial cells (η_21_). In this case, the lower threshold of η_me_ can indeed shift to a smaller value (Figure 2D), which makes it theoretically possible to switch the system from state II (low epithelial cells) to state I (high epithelial cells) at a fairly small η_me_ value.

In summary, our results for Model II suggest that one can switch the system from state II (low epithelial cells) to state I (high epithelial cells) is to increase both η_me_ and the effective conversion from M_2_ to M_1_.

**Figure 2.**
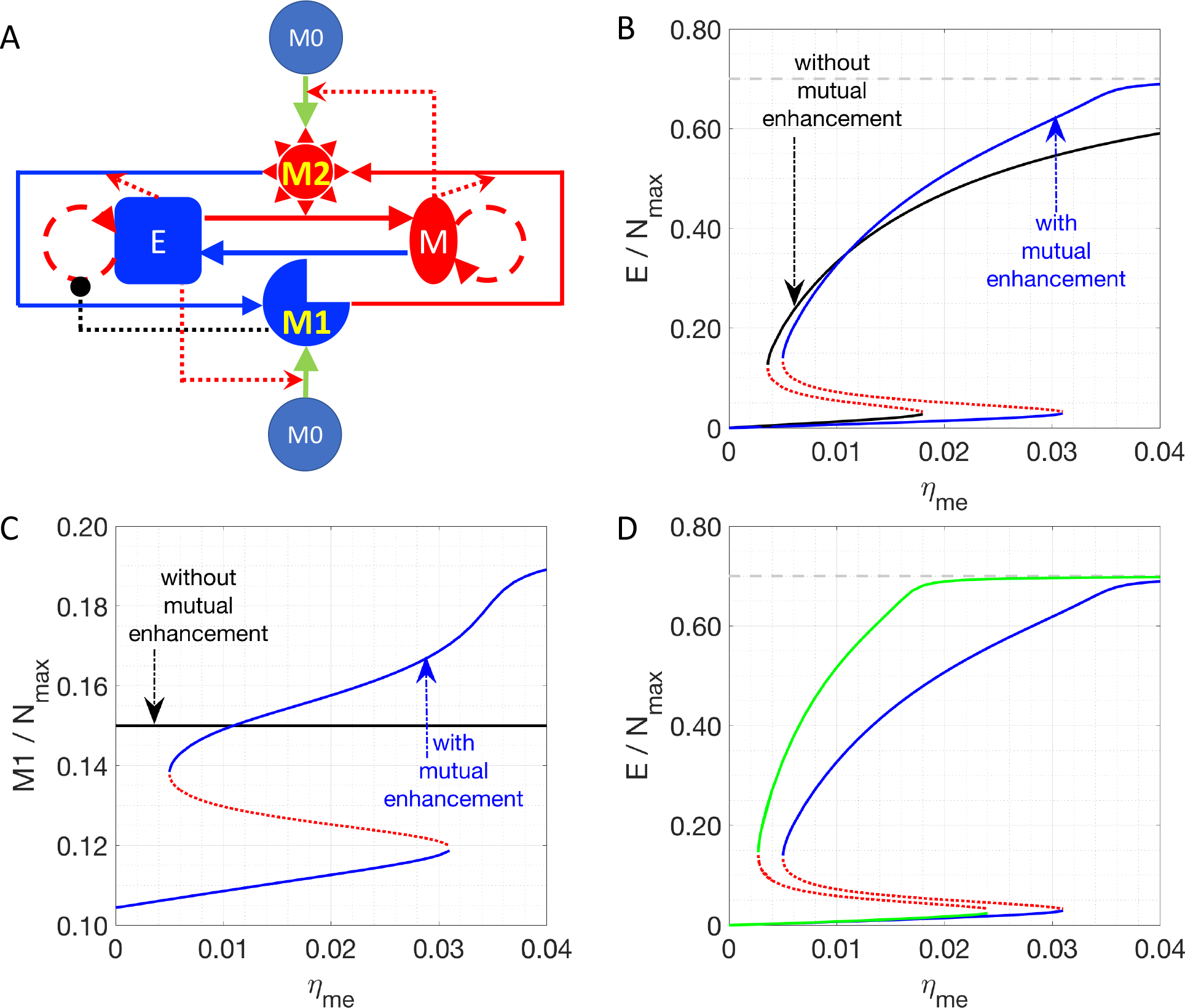
Effects of interconversion between M_1_ and M_2_ cells, mediated by cancer cells. A. Network of Model II. B and C. Steady states of the epithelial (B) or M_1_ (C) populations are plotted as a function of η_me_. Stable steady states are plotted in solid lines and unstable steady states are plotted in dotted lines. The key parameters in B and C are: η_12_=η_21_=0 (for black lines) or 1/72 h^−1^ (for blue lines), m=2, n=2, k=4, η^0^21η=^0^ =1/72 h^−1^. D. In this plot, the key parameters are: η_21_=1/72 h^−1^ (for blue lines) or 1/36 h^−1^ (for green lines), m=2, n=2, k=4, η^0^_21_=η^0^_12_=1/72 h^−1^, and η_12_=1/72 h^−1^. In B and D, the grey line is the maximum fraction of cancer cells (=0.7) as M_c_ is set as 0.3.

### Cell apoptosis-induced M1-to-M2 conversion leads to symmetry breaking in the cancer-immune interaction network

Finally, we have incorporated one other set of experimentally documented interactions, i.e. M_1_ macrophages can drive the apoptosis of epithelial cells, and factors released during cancer cell apoptosis can drive M_1_-to-M_2_ conversion (referred as Model III, Figure 3A, see Methods) (35). This interaction induces ‘symmetry breaking’ (46) in the system, as previously most of the interactions considered were of a ‘symmetric’ nature, i.e. M_1_ cells driving MET and M_2_ cells driving EMT, and epithelial cells driving M_1_ maturation while mesenchymal cells driving M_2_ maturation. With this new interaction, the system is now biased against epithelial cells, because a) epithelial but not mesenchymal cells, are killed by M_1_-macrophages, and b) their dead counterparts may convert M_1_ to M_2_ cells that can, in turn, convert some epithelial cells to mesenchymal ones (EMT).

Thus, in the parameter region investigated, there are 3 types of stable steady states, which are represented by solid blue, red and black lines, respectively. At small values of the control parameter η_me_ (rate of M_1_ macrophage assisted MET), there is only one type of stable steady state: the system is biased towards the mesenchymal-dominated state (solid blue curves in Figures 3B-3E), whereas the cancer-extinction state (E=M=0, dashed black curves in Figures 3B-3E) is unstable. As η_me_ increases and goes across a critical value η^a^_me_=0.0626 h^−1^), the extinction state (E=M=0) becomes stable as well as a new set of steady states emerges (red lines in Figures 3B-3E). Between η_me_=0.0663 h^−1^ and η_me_=0.0747 h^−1^, the steady states depicted as a solid red line (Figures 3B-3E) are stable; here both populations of epithelial and mesenchymal cancer cells are at a low level. This phenomenon can be understood in the following sense: at higher η_me_, proliferating mesenchymal cells are continuously being converted to epithelial cells, which will be killed by M_1_ macrophages. This effect brings down the overall fraction of cancer cells. Note that in this region, three types of stable steady states co-exist in the system for a given set of parameters. As η_me_ increases and goes across η^b^_me_=0.0747 h^−1^, the stable steady states in solid red lines disappears and the other two types of stable steady states co-existed (solid blue and black lines in Figures 3B-3E). As η_me_ further increases and goes across η^c^_me_=0.1532 h^−1^, there is only one stable steady state: cancer cells are necessarily eliminated from the system. We note in passing that the instability that drives the red solution unstable as η_me_ is lowered past 0.0663 h^−1^ is a Hopf bifurcation to an unstable limit cycle.

Furthermore, for the breast cancer cell line MDA-MB-231 used in Ref. (38), η_me_ is estimated to be around 1/120 h^−1^ (~0.0083 h^−1^). In order to explore the conditions for the absolute extinction of cancer cells around the estimated η_me_, we varied different parameters (such as λ_M_, η_em_ and η_21_) to get the corresponding η^c^_me_ where the extinction state is the only stable steady state of the system. We found that lowering the growth rate of mesenchymal cells (η_M_) can reduce η^c^_me_ to the nominal experimental value 1/120 h^−1^ (Figure 3F) whereas lowering down η_em_ or increasing η_21_ might not (Figure S5). Therefore, for therapeutic purposes, increasing η_me_ and decreasing η_M_ can be a promising combination to help to eliminate cancer cell populations.

**Figure 3.**
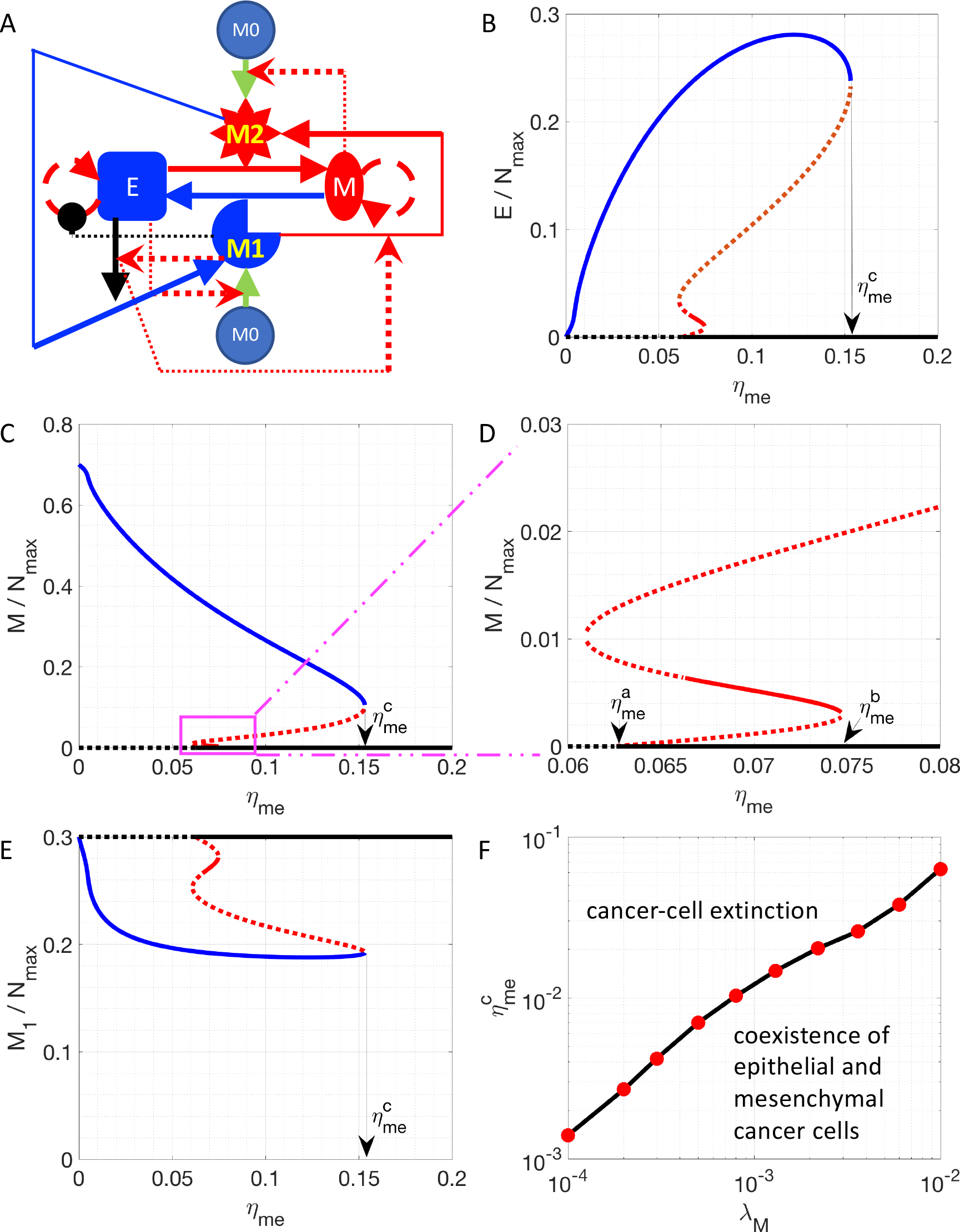
Steady state solutions for a model including the M_1_ to M_2_ conversion assisted by apoptotic epithelial cancer cells. A. Network of Model III. B-E. Steady states of the epithelial (B), mesenchymal (C and D) and M_1_ (E) populations are plotted as a function of ηme. D is a zoomed-in version of C. Stable steady states are plotted in solid lines and unstable steady states are plotted in dotted lines. The model-specific parameters are: β=1/36 h^−1^, β_c_=1/1200 h^−1^, η12η=_21_=1/72 h^−1^. When η_me_ is higher than a critical value η^a^_me_, the steady state E=M=0 becomes stable (D). When η_me_ is higher than a critical value η^c^_me_, the steady state E=M=0 is the only stable steady state of the system. F. For a given λ_M_, the critical value η^c^_me_ is plotted.

### EMT scores correlate with multiple genes upregulated in M2 macrophages

As a proof of principle for the predictions of our model, we investigated multiple TCGA datasets, using our previously developed EMT scoring metric (47). This metric quantifies the extent of EMT in a particular sample, and correspondingly assigns a score between 0 (fully epithelial) and 2 (fully mesenchymal). We calculated the correlation coefficients for EMT scores with various genes reported to be differently regulated in M_1_ or M_2_ macrophages relative to M_0_ macrophages. We observed that many genes upregulated in M2 and downregulated in M_1_ macrophages – ACTN1, FLRT2, MRC1, PTGS1, RHOJ, TMEM158 – correlated positively with the EMT scores (Table 1, first 6 rows) across multiple cancer types. The higher the EMT scores, the higher the levels of those genes, including the canonical M_2_ macrophage marker CD206 (MRC1). On the other hand, many genes upregulated in M_1_ macrophages but downregulated in M_2_ macrophages – FIIR, STAT1, RSAD2, TUBA4A and XAF1 – showed either a negative or an overall weak positive correlation with EMT scores (Table 1, last 5 rows). The only exception observed in this trend was that for ARHGP24 which correlated positively with EMT scores across cancer types. Together, these correlation results in multiple TCGA datasets offer a promising initial validation of our model predictions that a state dominated by epithelial cells typically has higher number of M_1_ macrophages, while the other state dominated by mesenchymal cells typically has higher number of M_2_ macrophages.

**Table 1:** Correlation coefficients of expression levels of specific genes with EMT scores, across many TCGA datasets. FC is the fold-change of gene expression levels in M_1_ or M_2_ macrophages relative to M_0_, which is measured in Ref. (48) for murine bone marrow derived macrophages polarized *in vitro*. Those shown in red indicated upregulated in M_2_, those in blue indicate upregulated in M_1_.

| Gene | FC<br>(M1/M0) | FC<br>(M2/M0) | Correlation<br>(Colorectal) | Correlation<br>(Lung) | Correlation<br>(Pancreatic) | Correlation<br>(Breast) |
| --- | --- | --- | --- | --- | --- | --- |
| ACTN1 | 0.46 | 2.37 | 0.32 | 0.41 | 0.32 | 0.13 |
| FLRT2 | 0.47 | 7.29 | 0.33 | 0.36 | 0.33 | 0.40 |
| MRC1 (CD206) | 0.02 | 2.82 | 0.21 | 0.31 | 0.21 | 0.28 |
| PTGS1 | 0.44 | 11.02 | 0.28 | 0.47 | 0.28 | 0.18 |
| RHOJ | 0.21 | 2.96 | 0.36 | 0.30 | 0.36 | 0.35 |
| TMEM158 | 0.37 | 2.99 | 0.24 | 0.31 | 0.24 | 0.14 |
| ARHGAP24 | 2.38 | 0.36 | 0.29 | 0.24 | 0.29 | 0.37 |
| FIIR | 3.43 | 0.41 | -0.12 | -0.44 | -0.12 | -0.19 |
| STAT1 | 2.27 | 0.27 | -0.02 | 0.29 | -0.02 | 0.04 |
| RSAD2 | 3.03 | 0.09 | 0.08 | 0.27 | 0.08 | 0.02 |
| TUBA4A | 2.18 | 0.33 | 0.11 | -0.12 | 0.11 | -0.01 |
| XAF1 | 2.09 | 0.25 | 0.05 | 0.27 | 0.05 | 0.05 |

## Discussion

The tumor microenvironment involves multiple cell types that interact among each other in diverse ways, thus giving rise to a complex adaptive ecological system (49–51). For such a system, mathematical models can help reveal the mechanisms underlying the emergent behavior and eventually aid in designing effective therapeutic strategies to modulate those aspects of the microenvironment that fuel disease aggressiveness (52–58).

Here, we focused on the interactions among macrophages of different polarizations (M_1_ and M_2_) and cancer cells with different phenotypes (epithelial and mesenchymal). Based on the literature, we focused on two types of models: with (Model II and III) or without (Model I) interconversion between M_1_ and M_2_ macrophages. All three models share a common feature: with a given set of parameters, multiple types of stable steady states can co-exist. More specifically, in Model I (without M_1_-M_2_ interconversion), with a given set of parameters, the system can converge to continuous range of states depending on the initial condition. However, these steady states belong to two categories: state I with a higher epithelial population and state II with a lower epithelial population. After the system reaches a steady state, perturbations applied only to cancer cell populations cannot drive the system out of the original steady state whereas perturbations on macrophage populations will drive the system out of the original steady state. However, the perturbation on macrophage populations might not drive the system out of the original category of steady states unless the perturbation is sufficiently strong. In Model II and III, with a given set of parameters, the system can again converge to two types of steady states depending on the initial condition: state I with a higher epithelial population and state II with a lower (or even zero for Model III) epithelial population. Now however, the states are discrete. After the system reaches a steady state, perturbations of any single cell population might not drive the system out of the original steady state. Therefore, in general, it is not easy to switch the cancer-macrophage system from a mesenchymal-and M_2_-dominated state to an epithelial-and M_1_-dominated state.

Mathematical approaches similar to ours may be useful in explaining, and even predicting, the efficacy of different therapeutic intervention(s) and their combinations. For example, various efforts have been made to switch populations of macrophages from M_2_-into M_1_-like (25), such as depletion of TAMs, reprogramming of TAMs toward M_1_-Like macrophages, inhibition of circulating monocyte recruitment into tumor, etc. However, it is unclear that how effective these types of strategies would be. Our modeling studies suggest that when we consider the interactions in Model III, the efforts on manipulating the M_1_-M_2_ interconversion might not be effective to eliminate cancer cells; whereas in Model II increasing the M_2_-to-M_1_ conversion rate can help the system to switch to the epithelial cancer cell and M_1_ macrophage dominated state, which is believed to be less aggressive. Therefore, the effective therapeutic strategies strongly depend on the type of interactions present between cancer cells and macrophages.

It is important to recognize that our model suffers from limitations. For instance: a) it does not consider spatial aspects of these interactions, for instance, mesenchymal cells may migrate and invade through the matrix, thus changing the interactions among the cells considered in our framework; b) it does not consider the effects of senescence on epithelial cell growth (59); and c) it considers EMT and macrophages polarization as a binary process, whereas emerging reports support the notion that in both cases there is likely to exist a spectrum of states/phenotypes (60,61). A more realistic model that can overcome the abovementioned assumptions and thus reflect the dynamics of tumor microenvironment more accurately can be used to help designing a more effective way to switch and stably maintain the system into a less aggressive state.

Despite the abovementioned limitations, our model can contribute to identifying key parameters of the system. For example, it suggests that to design an effective therapy to maintain the system in a M_1_-dominated and cancer-free steady state, not only the conversion rate from mesenchymal to epithelial cells should be significant, but also the growth rate of mesenchymal cells should be low enough. In other words, MET-inducing and cell-growth-suppressing can together synergistically restrict disease aggressiveness.

In summary, our results show that the interaction network between tumor cells and macrophages may lead to multi-stability in the network: one state dominated by epithelial and M_1_-like cells, the other dominated by mesenchymal and M_2_-like cells. We also identify that inducing MET and inhibiting cancer-cell growth might be much more efficient in ‘normalizing’ (1) the tumor microenvironment that can otherwise be engineered by cancer cells to support their growth (62) .

## Method

### Computational Models

According to in vitro experiments in literature, we construct three mathematical models. In Model I, factors secreted by epithelial (mesenchymal) cancer cells will polarize monocytes into M_1_-like (M_2_-like) macrophages. M_1_-like macrophages will induce senescence of epithelial cancer cells and convert mesenchymal cancer cells to epithelial ones. On the other hand, M_2_-like macrophages will convert epithelial cancer cells to mesenchymal ones. In addition, a mesenchymal cancer cell can secrete soluble factors to help maintain the mesenchymal state of itself and its neighbors, whereas an epithelial cancer cell can maintain the epithelial state through being in contact with its neighboring epithelial cancer cells. There is assumed to be no cell growth or death for macrophages and there is a maximum ‘carrying capacity’ of cells in the system. The figure that illustrate this model is in Fig. 1A. The equations to describe this system is as follows:

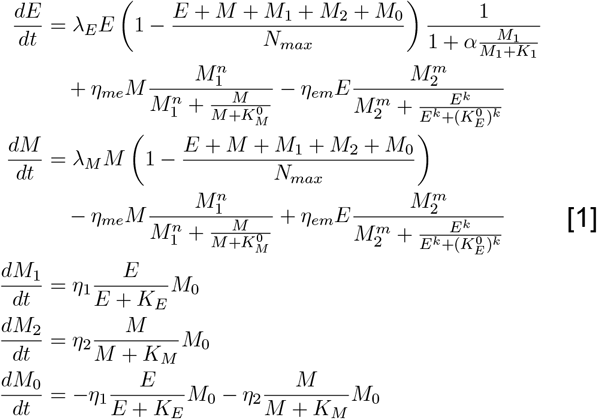

The description and the value of each parameter are given in Table 2.

In Model II, interconversions between M_1_ and M_2_ macrophages are included. Furthermore, the interconversions can be enhanced by corresponding cancer cells. The figure that illustrate this model is in Fig. 2A. The equation to describe this system is as follows:

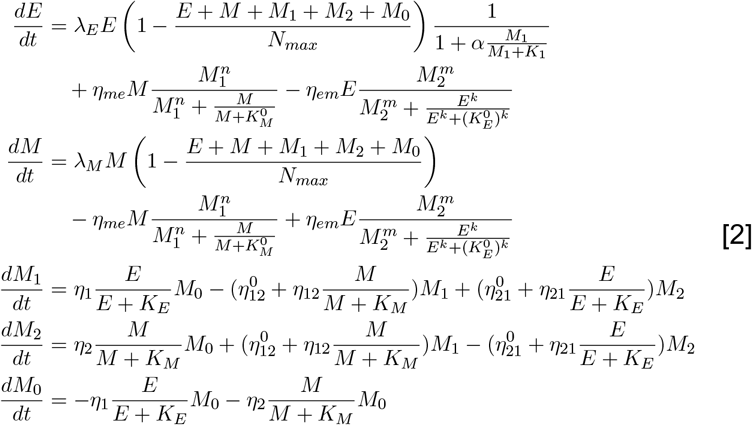

In the third model, additional interactions were introduced as illustrated in Fig. 3A. M_1_-like macrophages can induce apoptosis of epithelial cancer cells and factors released by apoptotic cancer cells can convert M_1_-like macrophages into M_2_-like macrophages. In order to restore the symmetry of the system, we further consider the therapeutic interaction: M_2_-like macrophages can be re-polarized back to M_1_-like macrophage by Type 1 T helper cells, IL-12, IFN-gamma, etc. The equation to describe this system is as follows:

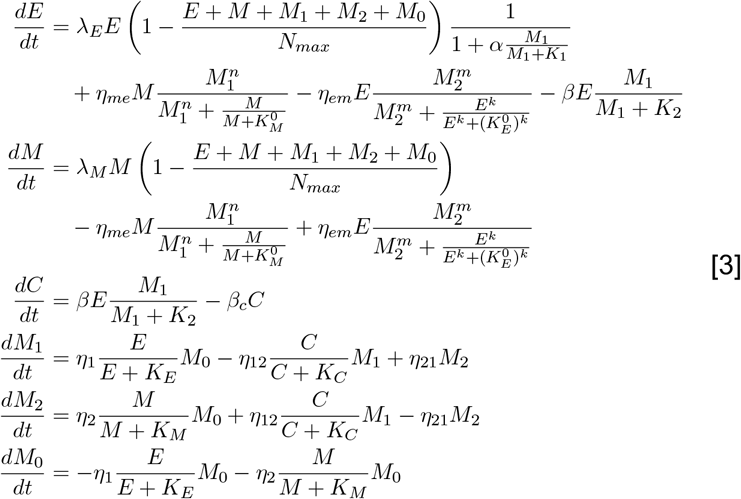

The description and the corresponding values of additional parameters are given in Table 2.

**Table 2:**
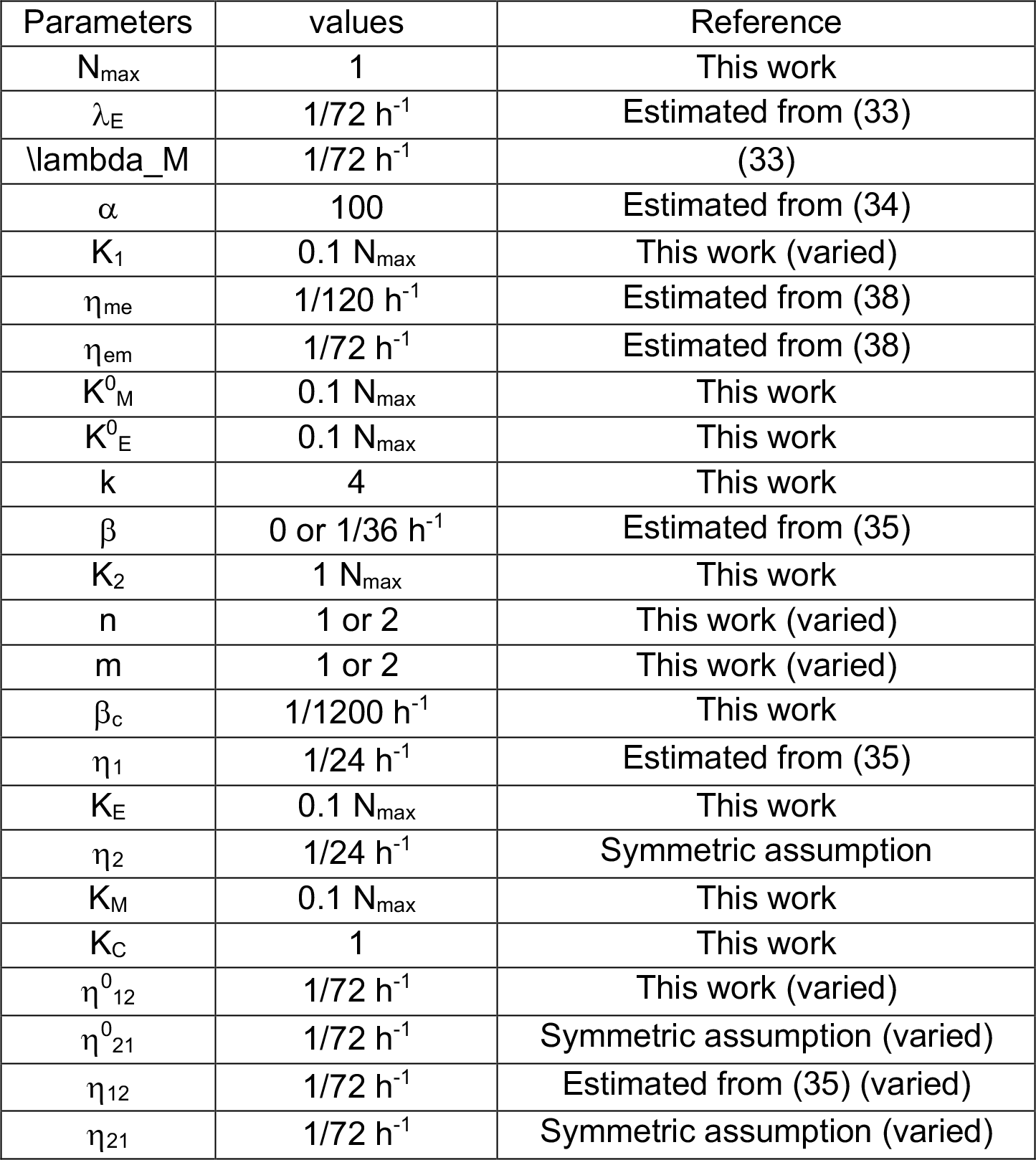
Parameters used in our models.

| Parameters | values | Reference |
| --- | --- | --- |
| $N_{\max}$ | 1 | This work |
| $\lambda_E$ | $1/72 \text{ h}^{-1}$ | Estimated from (33) |
| $\lambda_M$ | $1/72 \text{ h}^{-1}$ | (33) |
| $\alpha$ | 100 | Estimated from (34) |
| $K_1$ | $0.1 N_{\max}$ | This work (varied) |
| $\eta_{me}$ | $1/120 \text{ h}^{-1}$ | Estimated from (38) |
| $\eta_{em}$ | $1/72 \text{ h}^{-1}$ | Estimated from (38) |
| $K_M^0$ | $0.1 N_{\max}$ | This work |
| $K_E^0$ | $0.1 N_{\max}$ | This work |
| $k$ | 4 | This work |
| $\beta$ | 0 or $1/36 \text{ h}^{-1}$ | Estimated from (35) |
| $K_2$ | $1 N_{\max}$ | This work |
| $n$ | 1 or 2 | This work (varied) |
| $m$ | 1 or 2 | This work (varied) |
| $\beta_c$ | $1/1200 \text{ h}^{-1}$ | This work |
| $\eta_1$ | $1/24 \text{ h}^{-1}$ | Estimated from (35) |
| $K_E$ | $0.1 N_{\max}$ | This work |
| $\eta_2$ | $1/24 \text{ h}^{-1}$ | Symmetric assumption |
| $K_M$ | $0.1 N_{\max}$ | This work |
| $K_C$ | 1 | This work |
| $\eta_{12}^0$ | $1/72 \text{ h}^{-1}$ | This work (varied) |
| $\eta_{21}^0$ | $1/72 \text{ h}^{-1}$ | Symmetric assumption (varied) |
| $\eta_{12}$ | $1/72 \text{ h}^{-1}$ | Estimated from (35) (varied) |
| $\eta_{21}$ | $1/72 \text{ h}^{-1}$ | Symmetric assumption (varied) |

#### Correlation analysis

For a fixed dataset, linear regression was performed for each gene of interest against the predicted EMT score(47). The linear correlation coefficient was recorded in each case. We performed statistical analysis under the null hypothesis of zero correlation between gene expression fold-change and EMT score and recorded the corresponding p-values at significance level α=0.05. All datasets were obtained from the R2: Genomics Analysis and Visualization Platform (http://r2.amc.nl).

## Acknowledgements

This work was sponsored by the National Science Foundation NSF grant PHY-1427654 (Center for Theoretical Biological Physics). XL was supported by Stand Up to Cancer and The V Foundation. MKJ was also supported by a training fellowship from Gulf Coast Consortium as computational cancer biology training grant (CPRIT RP1705593). JTG was supported by National Cancer Institute of NIH (F30CA213878). KJP is supported by National Cancer Institute NCI grant CA093900. HL is also a Cancer Prevention and Research Institute of Texas Scholar in Cancer Research of the State of Texas at Rice University.

